# ETP-Specific Knockout Mice Reveal Endotrophin as a Key Regulator of Kidney Fibrosis in Ischemia-Reperfusion Injury Models

**DOI:** 10.1101/2025.02.20.639288

**Authors:** Dae-Seok Kim, Jan-Bernd Funcke, Shiuhwei Chen, Kyounghee Min, Toshiharu Onodera, Min Kim, Joselin Velasco, Megan Virostek, Katarzyna Walendzik, Philipp E Scherer

## Abstract

Endotrophin (ETP), a cleavage product of the C5 domain of collagen VI α3 (COL6A3), plays a crucial role in extracellular matrix remodeling. Previously established *Col6a3* knockout (KO) mouse models primarily reflect the consequences of COL6A3 loss rather than the specific effects of ETP depletion, making it challenging to directly assess ETP’s function. These models either disrupt COL6A3 along with ETP production or express functionally defective COL6A3 while maintaining ETP production. To address this limitation, we developed and validated a novel ETP knockout (ETPKO) mouse model that selectively ablates ETP while preserving *Col6a3* expression. To generate the ETPKO model, we introduced *lox2272* sites and a fluorescent *mCherry-CAAX* reporter into the *Col6a3* locus, ensuring that ETP expression is turned off and reporter expression is turned on upon Cre-mediated recombination. Crossing the *Col6a3-Etp+mCherry-CAAX* mouse line with CMV-Cre mice yielded ETPKO mice, in which successful ETP deletion was confirmed by genomic DNA sequencing and *mCherry* expression. Using this model, we investigated ETP’s role in kidney fibrosis. ETPKO mice subjected to unilateral or bilateral renal ischemia-reperfusion injury (IRI) exhibited complete *Etp* mRNA ablation with only a partial reduction in *Col6a3* mRNA. Notably, ETP depletion significantly attenuated fibrosis progression, demonstrating its critical role in the pathogenesis of kidney fibrosis. The ETPKO mouse model provides a targeted and specific approach for studying ETP function independently of *Col6a3* expression. These findings establish ETP as a key driver of fibrosis and position ETPKO mice as a valuable tool for elucidating ETP-mediated mechanisms in preclinical disease models.

## INTRODUCTION

Collagen type VI (COL6), a critical component of the extracellular matrix, plays a crucial role in maintaining tissue structure and integrity ^1–3^. Composed of three chains (α1, α2, and α3) encoded by *Col6a1, Col6a2,* and *Col6a3,* COL6 forms tetramers that aggregate into microfibrils within the extracellular space ^2,3^. Endotrophin (ETP), a cleavage product of the COL6A3 chain, is derived from its C5 domain, which is proteolytically cleaved from COL6 fibrils immediately after secretion ^4–7^. This cleavage and the resulting separation suggest that ETP has a functional role distinct from COL6A3. Various *Col6a3* mutant mouse models have been developed, differing significantly in their capacity to produce functional ETP. While these models have been used to study the biology of ETP, they clearly lack specificity due to manifest defects in COL6 function. Each model exhibits fundamentally different impairments in COL6 function and potentially distinct effects on ETP production. ^8,9^. The first model carries a hypomorphic *Col6a3* allele (*Col6a3hm*), which leads to a near-complete absence of *Col6a3* transcription, disrupted intracellular COL6 dimer and tetramer assembly and secretion, and associated muscle and tendon defects ^8^. The second model carries a disrupted *Col6a3* allele (*Col6a3d16*), which causes an in-frame deletion of 18 amino acids at the N-terminus of the triple-helical domain of the COL6A3 parent chain ^9^. While heterozygous *Col6a3+/d16* and homozygous *Col6a3d16/d16* mice exhibit normal intracellular COL6 dimer and tetramer assembly and secretion, they display disrupted extracellular microfibril formation along with muscle and tendon defects. Importantly, whereas the *Col6a3hm* allele is expected to decrease ETP production, the *Col6a3d16* allele likely maintains ETP production. These findings suggest that the COL6A3 parent form is critical for muscle and tendon integrity, whereas ETP may not play a significant role in this context. Nonetheless, neither model provides definitive insight into the contribution of ETP to the studied biological processes. To address these shortcomings, we now developed a novel *Col6a3-Etp+mCherry-CAAX* mouse line using CRISPR/Cas9 genome editing to selectively modify the *Col6a3* locus upstream of the endogenous endotrophin (ETP)-coding sequence. In this model, ETP can be specifically deleted through Cre-mediated recombination, enabling functional studies of ETP without affecting the COL6A3 parent chain. ETP has been identified as a critical mediator of fibrosis, driving disease progression through fibrotic processes and inflammation and serving as a key factor in the pathophysiology of fibro-inflammatory disorders ^10–12^. It stimulates fibroblasts to upregulate collagen I (COL1) synthesis, a key component of fibrosis, and elevated ETP levels have been associated with fibrosis and adverse outcomes in heart failure with preserved ejection fraction (HFpEF) ^13^. Additionally, ETP is strongly associated with fibrosis-related conditions such as chronic kidney disease (CKD), where high ETP levels have been independently linked to increased mortality risk, underscoring its role in disease progression and prognosis ^14^. Recently, our group demonstrated that targeting ETP with neutralizing antibodies reduces renal fibrosis and improves renal function in a mouse model of CKD, further supporting ETP’s role in chronic fibro-inflammatory kidney diseases ^15^. Collectively, these findings highlight the role of ETP as both a biomarker and a contributing factor in fibrotic conditions, providing a strong rationale for investigating its specific role in fibrosis independently of its parent gene, *Col6a3.* Unilateral or bilateral renal IRI is a well-established model for studying the progression of acute kidney injury to fibrosis ^16–18^. Here, we demonstrate that ETP is a critical driver of kidney fibrosis independent of COL6A3. Using our novel ETP-specific knockout (ETPKO) mouse model, we show that ETP depletion significantly reduces fibrotic protein expression and mitigates kidney fibrosis under ischemia-reperfusion injury conditions. These findings establish ETP as a key mediator of fibrosis and validate ETPKO mice as a robust tool for studying fibrotic processes.

## MATERIALS AND METHODS

### Study approval

All animal experimental protocols, including those for mouse use and euthanasia, were reviewed and approved by the Institutional Animal Care and Use Committee (IACUC) of the University of Texas Southwestern (UTSW) Medical Center under animal protocol numbers 2024-103554, 2024-103545-G, and 2024-103559-G.

### Generation of the Col6a3-Etp+mCherry-CAAX line

*Col6a3-Etp+mCherry-CAAX* mice were generated using CRISPR/Cas9-based genome editing. A guide RNA sequence (CAGTTCAACCATCAACCTCA-[TGG]) was designed *in silico* using CRISPOR (www.crispor.tefor.net) to target the mouse *Col6a3* open reading frame (ORF) upstream of the start of the endogenous endotrophin (ETP)-coding sequence, which naturally spans the last three exons of the gene. A donor plasmid containing a 350 bp left homology arm, a first in-frame lox2272 site, a copy of the full ETP-coding sequence with a stop codon, a second out-of-frame lox2272 site, a P2A-mCherry-CAAX-coding sequence with a stop codon, a copy of the full *Col6a3* 3’UTR, a rabbit β-globin poly(A) signal, and a 350 bp right homology arm was synthesized by Genewiz (annotated sequence provided in the supplement). This donor plasmid was used to prepare a linear, single-stranded repair construct using the Guide-it Long ssDNA Production System v2 (TaKaRa, #632666) with the following primers: 5’-TGGGTACTTAGGCTACACCCT G-3’ and 5’-GACCACAAGTCAACCCTAGCC-3’. An Alt-R CRISPR-Cas9 crRNA guide, tracrRNA, and Cas9 protein (all from IDT) were mixed with the repair construct and used for pronuclear injection into fertilized C57BL/6J eggs by the UTSW Transgenic Core. Injected eggs were transplanted into foster mothers, and the obtained offspring were screened for site-specific integration of the transgene by standard PCR. Candidate mice with apparent site-specific integration of the transgene were crossed to C57BL/6J mice. The offspring from these crosses were genotyped by PCR, and the full sequence of the integrated transgene, as well as the upstream and downstream regions, was verified by Sanger sequencing (Azenta/Genewiz). Sequence alignments were performed in SnapGene (version 8.0.2; GSL Biotech LLC). Fully sequence-verified mice were crossed to C57BL/6J mice for at least two more generations. We used the following primers for routine genotyping of the transgene: G336 5’- TCTTCAGGCAGCACACCGAG-3’; G337 5’-TCACCATAGGACCGGGGTTTT-3’; G338 5’- CTGAGGACCCCTTTGGAACTG-3’. These primers detect the modified (405 bp), non-modified (241 bp), and knockout (161 bp) alleles. For further validation of recombination, the PCR products of the modified and knockout alleles were isolated using the Monarch PCR & DNA Cleanup Kit (New England Biolabs # T1030L) and analyzed by Sanger sequencing (Azenta/Genewiz). Before Cre recombination, the modified *Col6a3-Etp+mCherry-CAAX* locus produces an mRNA that encodes the COL6A3 chain, a short 12 amino acid ‘ITSYRILYTKLS’ linker derived from the in-frame lox2272 site, and ETP. This mRNA possesses an untranslated region that includes the second out-of-frame lox2272 site, P2A-mCherry-CAAX-coding sequence, *Col6a3* 3’UTR, and a poly(A) tail. Importantly, the last two exons of the Col6a3 gene that encompass most of the endogenous ETP-coding sequence do not contribute to the *Col6a3-Etp+mCherry-CAAX* mRNA because transcription of the modified locus terminates upstream at the introduced rabbit β-globin poly(A) signal. Cre-mediated recombination of the lox2272 sites removes the ETP-coding sequence and brings the P2A-mCherry-CAAX-coding sequence in-frame. The resulting mRNA encodes the COL6A3 chain, a ‘ITSYRILYTKLS-GSG-ATNFSLLKQAGDVEENPG*P’* stretch derived from the lox2272 site and P2A peptide, and mCherry-CAAX, followed by an untranslated region that includes the *Col6a3* 3’UTR and a poly(A) tail. The P2A peptide causes ribosomal skipping during translation, resulting in the separation of the membrane-targeted red fluorescent mCherry-CAAX protein from the remainder of the COL6A3 chain (note the indicated ‘G*P’ cleavage site).

### Mouse maintenance

All mice used in this study, including littermate-controlled males, were maintained on a pure C57BL/6 genetic background. Mice were housed under barrier conditions on a 12-hour light/12-dark cycle in a temperature-controlled environment (22°C) with and libitum access to autoclaved water and diet. Cages were changed every other week, and constant veterinary supervision was provided. Diets used in this study include regular chow diet (LabDiet #5058).

### Genotyping PCR

Genotyping PCR assays were performed as described previously ^19^. Briefly, a small piece of the mouse tail tip was incubated in 50 mM NaOH at 95°C for 1.5 hours and neutralized with 10% _V/V_ 1M Tris-HCl (pH 8.0). The supernatant was used as a PCR template with primers listed in Table S1. PCR products were analyzed on a 1–2% agarose gel stained with ethidium bromide.

### Unilateral kidney ischemia-reperfusion model

The unilateral kidney ischemia-reperfusion model followed the protocol from previous studies ^16,17,20^. A mixture of ketamine (25 mg/ml) and xylazine (2.5 mg/ml) was administered via intraperitoneal injection to anesthetize the mouse. The anesthetized mouse was placed on an infrared warming pad (Kent Scientific Corporation) to maintain body temperature within 36–37 °C. To prevent dryness during anesthesia, Puralube® Vet Ointment was applied to the eyes. The hair around 1 cm lateral to the spine below the 13th rib, was shaved, and the skin was cleaned three times using 75% ethanol followed by povidone-iodine. A small incision was made using scissors, and the skin was bluntly separated from the peritoneum. The intestines were gently displaced toward the right side of the abdominal cavity using sterile, saline-moistened cotton swabs. The renal pedicles were exposed by carefully separating the fascia and adipose tissue using forceps. Renal ischemia was induced by applying a micro clip to the renal artery and vein, with successful ischemia visually confirmed by the gradual and uniform darkening of the kidney. After the designated ischemia duration, the clip was removed, and successful reperfusion was confirmed by the rapid color change of the kidney from dark red to dark pink. Finally, the peritoneum and skin were closed using Vicryl 5-0 sutures and clips, respectively.

### Tissue preparation for histological analysis

Mice were euthanized *via* cervical dislocation after isoflurane anesthesia. Tissues were fixed in 10% formalin for 24 hours at room temperature, stored with 50% ethanol, and embedded in paraffin and cut into 5 μm sections for histological analysis.

### Histopathological analysis

For histopathological phenotyping, two age-matched (10-week-old) mice of each gender per genotype were submitted to the UTSW ARC Diagnostic Lab. The mice were euthanized, and H&E-stained slides were prepared from multiple tissues. Each slide was reviewed by the ARC Diagnostic Lab to evaluate for unusual phenotypes.

### Immunostaining

Immunostaining assays were performed as described previously ^19^. Briefly, paraffin sections (5 μm) were deparaffinized, subjected to antigen retrieval, and blocked in 10% goat serum. Slides were incubated overnight at 4°C with primary antibodies in 5% BSA, including ETP, a-SMA COL6, mCherry or RFP, followed by goat-derived Alexa Fluor-labeled secondary antibodies for 1 hour at room temperature. After washing, slides were mounted with DAPI-containing medium and imaged using a Zeiss LSM880 confocal microscope provided by the UTSW Quantitative Light Microscopy Core Facility. Image analysis and quantification were performed using FIJI/ImageJ.

### H&E and picrosirius red staining

For H&E staining, paraffin-embedded sections were deparaffinized, rehydrated, and stained using an H&E Staining Kit (Abcam #ab245880). For picrosirius red staining, paraffin-embedded sections were deparaffinized, rehydrated, and sequentially stained with Weigert’s hematoxylin, phosphomolybdic acid, and picrosirius red, with rinses performed after each staining step. Sections were then rinsed in 0.1 N hydrochloric acid and 0.5% acetic acid, dehydrated in ethanol, and prepared for imaging. The entire kidney image was acquired using a NanoZoomer 2.0 HT (Hamamatsu) provided by the UTSW Whole Brain Microscopy Facility. Image analysis was performed using NDP.view 2.

### Preparation of whole cell extracts from tissues and immunoblotting

Frozen tissues were pulverized and resuspended in RIPA Buffer (Pierce) using a glass douncer on ice. The mixture was incubated at 4°C for 20 minutes with gentle mixing, followed by centrifugation to remove insoluble material. The supernatant was collected as the soluble extract, and protein concentrations were determined using a BCA Protein Assay (Pierce). For Western blotting, 20 μg of protein was separated on a 4-12% gradient SDS-PAGE gel (Invitrogen) and transferred to a nitrocellulose membrane (BioRad). Membranes were blocked in 5% nonfat dry milk and incubated overnight at 4°C with primary antibodies, including ETP, a-SMA, COL1, COL6, mCherry or RFP, and GAPDH. After washing, HRP-conjugated secondary antibodies (ThermoFisher) were applied, and protein signals were detected using chemiluminescence with the iBright 1500.

### Quantitative PCR (qPCR)

Total RNA was extracted using RNeasy Mini Kit, Trizol, and EZ-10 DNAaway RNA Miniprep Kit, followed by cDNA synthesis with PrimeScript™ RT Master Mix. qPCR was performed using PowerUp SYBR Green Master Mix on a QuantStudio 6 Flex System. Primer sequences are listed in Table S2.

### Statistics

Statistical analyses were performed using Prism (version 10.4.1; GraphPad), applying two-tailed Student’s t-tests for pairwise comparison and one-way ANOVA. Statistical significance was set at p < 0.05.

## RESULTS

### Generation and validation of whole body ETP knockout mice

To investigate the physiological and pathophysiological functions of ETP, we generated systemic ETP knockout (ETPKO) mice. To this end, we first generated and sequence-verified a Cre-dependent *Col6a3-Etp+mCherry-CAAX* knockin line and then crossed this line with a germline Cre recombinase driver (CMV-Cre) (Fig. 1a and Supplementary Fig. 1a). Genotyping validation for the knockin, wild-type (WT), and knockout (ETPKO) alleles was performed using primers spanning the targeted genomic regions (Fig. 1b). The Col6a3-Etp-mCherry-CAAX knockin line was designed to allow Cre recombination to selectively turn off ETP expression while simultaneously turning on the expression of a membrane-bound red fluorescent protein reporter (mCherry-CAAX). Following a single-generation cross with CMV-Cre, we selected mice lacking the CMV-Cre allele to establish a colony for our experiments (Fig. 1c). For further validation, we also utilized Sanger sequencing to confirm the successful deletion of the ETP-encoding sequence in ETPKO mice (Supplementary Fig. 1b). Consistent with this finding, immunoblot (Fig. 1d) and immunofluorescence (IF) analyses (Fig. 1e) of kidney tissues demonstrated mCherry expression exclusively in the ETPKO mice but not in WT littermate mice, confirming that the designed switch to reporter expression following ETP deletion worked as intended. Furthermore, RT-qPCR analysis across multiple tissues revealed complete ablation of *Etp* mRNA expression in the ETPKO mice, while its parental transcript, *Col6a3*, exhibited only a mild but non-significant reduction in expression compared to WT littermates (Fig. 1f and Supplementary Fig.1c). Importantly, *Col6a3* expression in ETPKO mice remained sufficient to rule out significant functional contributions from its depletion, with the observed mild reduction potentially arising as a secondary effect of ETP elimination which can act as a feed-forward profibrotic factor. To assess whether ETPKO mice develop normally, we conducted a histopathological analysis of multiple H&E-stained tissues, evaluated by UTSW ARC Diagnostic Lab, which revealed no apparent morphological changes compared to WT littermates (Supplementary Fig. 2a). Body weights and body condition scores were also comparable between WT and ETPKO mice (Supplementary Fig. 2b, c). Muscle weights showed no significant differences between WT and ETPKO mice (Supplementary Fig. 3a). Furthermore, H&E staining of gastrocnemius and diaphragm muscles indicated no overt phenotypic abnormalities (Supplementary Fig. 3b). In summary, we successfully generated a new transgenic ETPKO mouse model that lacks *Etp* mRNA expression altogether while displaying only a mild reduction of *Col6a3* mRNA expression and no overt phenotype in the absence of a specific challenge. The ETPKO model thus enables us for the first time to study the functional consequences of ETP deletion without confounding effects of altered COL6A3 function.

**Fig. 1.**
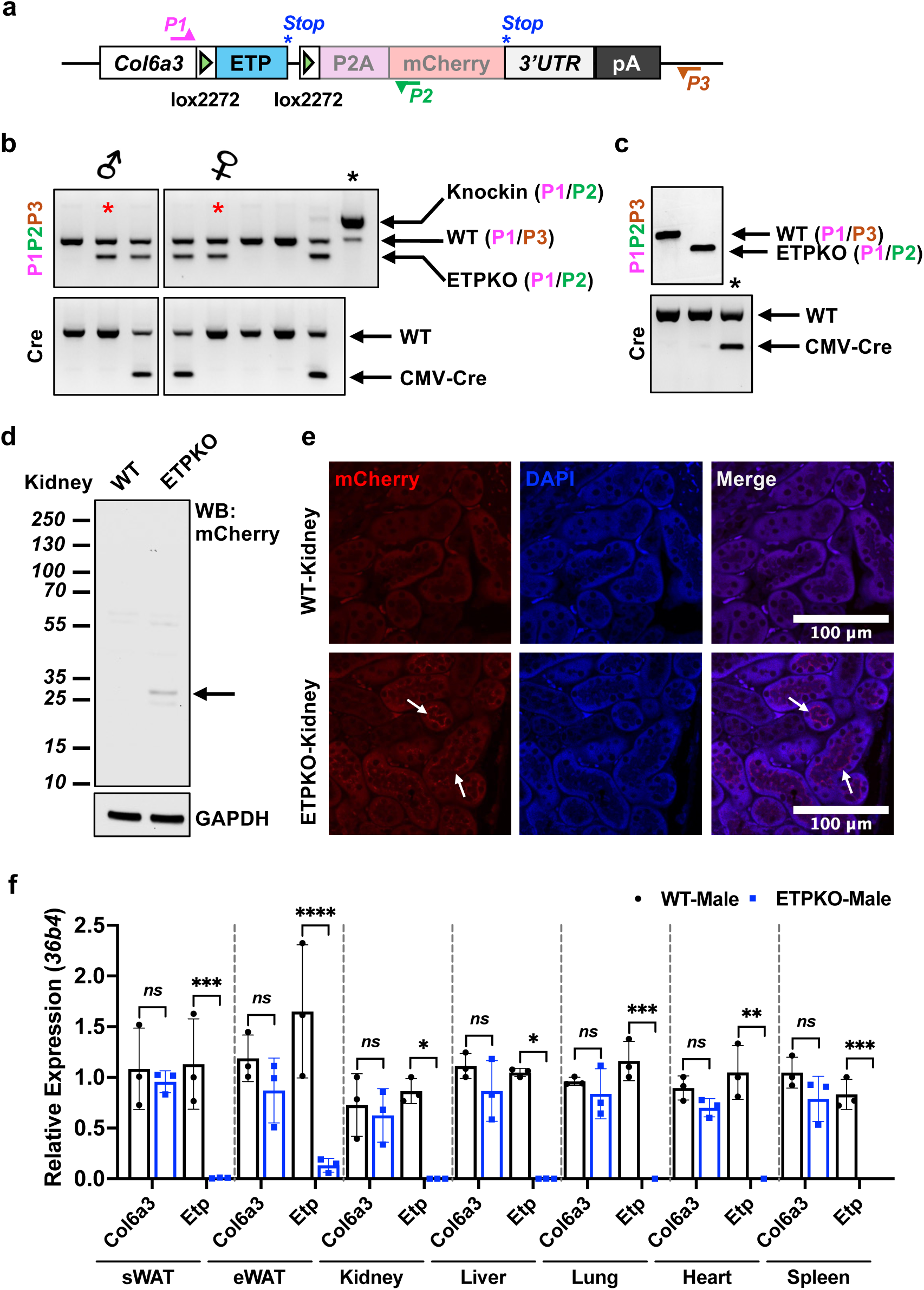
Generation and validation of whole body ETP knockout mice. **a** Schematic representation of the *Col6a3-ETP+mCherry-CAAX* allele before Cre-mediated recombination. The binding sites of genotyping primers P1 (pink), P2 (green), and P3 (orange) are indicated. The stop codons of the ETP and P2A-mCherry-CAAX reading frames are shown as asterisks (blue). **b-c** Genotyping validation for WT and ETPKO mice. The red asterisk indicates breeder mice, and the black asterisk indicates the positive control. **d** Immunoblot analysis of mCherry and GAPDH in kidney tissues from WT and ETPKO mice. **e** Representative immunofluorescence (IF) staining of mCherry in the kidney, reporting active COL6A3 expression (white arrows) (n = 3 male mice/group). **f** *Col6a3* and *Etp* mRNA expression, normalized to *36b4*. Data are presented as the mean± SEM (n = 3 male mice/group) and were analyzed by two-tailed Student’s t-tests. *, p<0.05; ** p<0.01; *** p<0.001.

### Induction of ETP expression following kidney ischemia-reperfusion injury and reduction of local fibrosis upon ETP depletion

ETP is widely recognized for its role in promoting fibrosis ^12,13,21^. To investigate its involvement in kidney fibrosis, we utilized a unilateral renal ischemia reperfusion injury (IRI) model, a well-established approach for inducing acute kidney injury and subsequent fibrosis ^16^. A schematic representation of the experimental design illustrates the procedure, which involves a 30-minute clamp of the renal artery followed by a 7-day postoperative recovery period (Fig. 2a). Post-IRI kidneys exhibited a visibly darker coloration compared to pre-IRI kidneys, confirming the successful induction of ischemia-reperfusion injury (Fig. 2b). Immunoblot analysis of kidney tissues revealed mCherry expression exclusively in ETPKO mice, confirming effective deletion of ETP. Notably, mCherry expression was highly upregulated in ETPKO-IRI kidneys, suggesting that IRI robustly induces *Col6a3* and, consequently, ETP expression (Fig. 2c). Histological analysis following H&E staining revealed severe pathological changes in WT-IRI kidneys, including a disruption of the tubular architecture and severe cyst formation. In contrast, ETPKO-IRI kidneys partially displayed preserved tubular architecture and reduced cyst formation, with notable dilation of the urine duct compared to WT-IRI kidneys (Fig. 2d). IF staining further demonstrated highly upregulated ETP expression in WR-IRI kidney of WT mice, while no expression was observed in WT-sham, ETPKO -sham, or ETPKO-IRI kidneys (Fig. 2e, f). The kidneys in each of these groups exhibited some non-specific signals at the boundary due to the boundary effects in immunostaining. We further assessed fibrosis using picrosirius red staining, which revealed extensive fibrosis in WT-IRI kidneys. Remarkably, fibrosis was significantly reduced in ETPKO-IRI kidneys, indicating that the absence of ETP impedes the fibrotic process (Fig. 2g). Collectively, these findings demonstrate that IRI robustly induces ETP expression in WT kidneys, while the lack of ETP in ETPKO mice significantly reduces kidney fibrosis, highlighting a critical role for ETP in the development of kidney fibrosis.

**Fig. 2.**
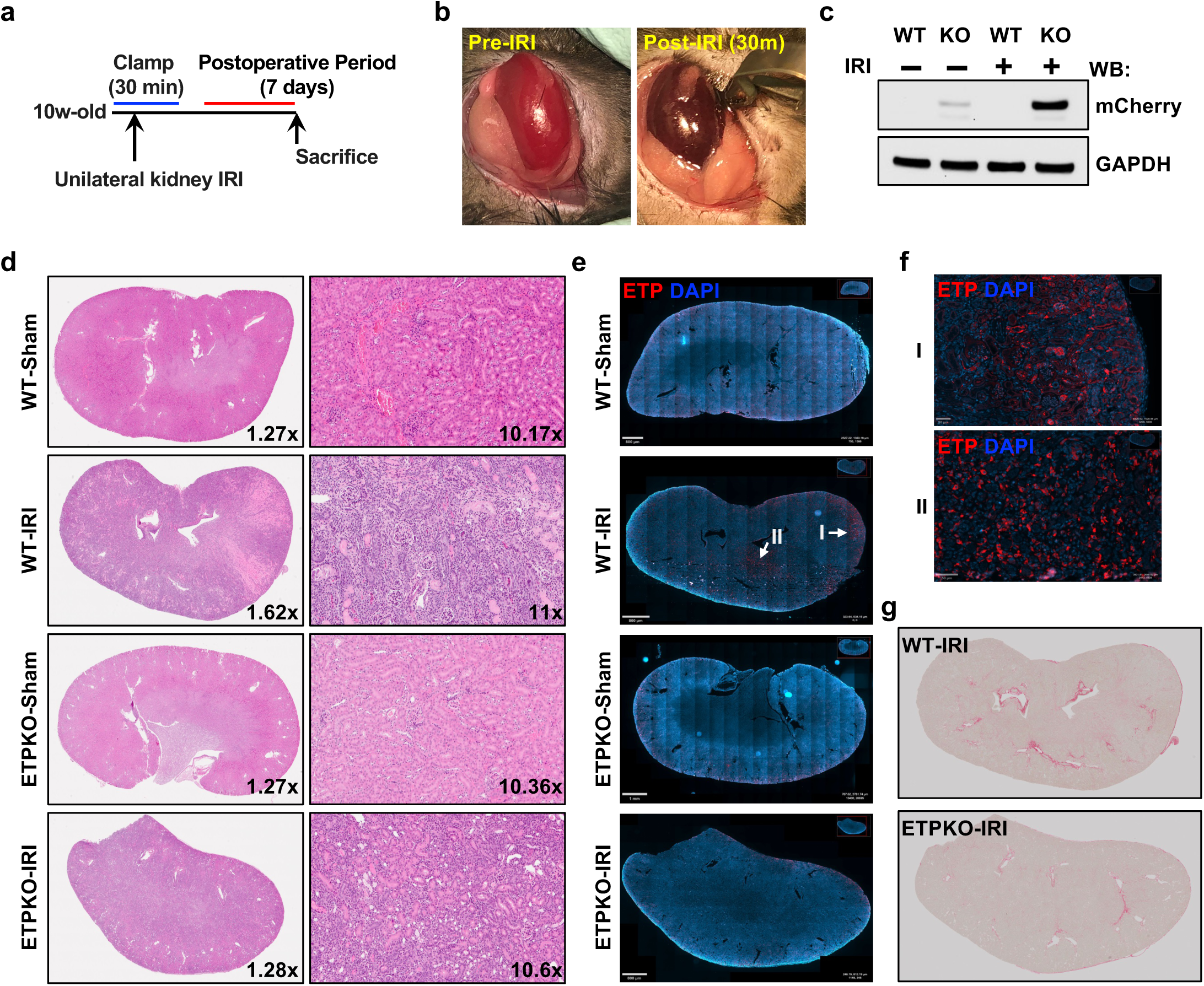
Induction of ETP expression following kidney ischemia-reperfusion injury and reduction of local fibrosis upon ETP depletion. **a** Schematic representation of the experimental design for the unilateral kidney ischemia-reperfusion injury (IRI) model. The procedure involves a 30-minute clamp of the renal artery, followed by a post-operative period of 7 days. **b** Images of the kidney pre- and post-IRI procedure. **c** Immunoblot analysis of mCherry and GAPDH in kidney tissues from WT and ETPKO mice under sham and post-IRI conditions **d** H&E staining showing the entire kidney and magnified regions for each group. **e** Representative IF staining of ETP across the whole kidney. **f** Magnified IF staining highlighting enriched ETP expression in the post-IRI kidney of WT mice (panel e). **g** Picrosirius red staining showing fibrosis in the entire kidney for WT-IRI and ETP KO-IRI groups.

### ETP depletion reduces local fibrotic gene mRNA and protein expression following kidney ischemia-reperfusion injury

To further delineate the role of ETP in regulating fibrotic gene expression, we extended the unilateral kidney IRI model by introducing a later post-operative recovery time point, specifically 12 days, following a 30-minute clamping of the renal artery (Fig. 3a). On post-operative recovery day 7, both WT-IRI and ETPKO-IRI kidneys exhibited a brighter appearance compared to sham kidneys (Fig. 3b, left panel). By post-operative recovery day 12, WT-IRI kidneys appeared slightly smaller than ETPKO-IRI kidneys (Fig. 3b, right panel), suggesting a potential impact of ETP on kidney structure during injury resolution. As observed in our previous analysis (Fig. 2), ETP was highly upregulated in WT mice following IRI. To confirm this, we analyzed *Col6a3* and *Etp* mRNA expression at days 7 and 12 post-IRI. On day 7, *Etp* mRNA was significantly upregulated in WT-IRI kidneys but was undetectable in ETPKO-IRI kidneys (Fig. 3c). Similarly, *Col6a3* mRNA was highly expressed in WT-IRI kidneys but significantly reduced in ETPKO-IRI kidneys (Fig. 3c). Interestingly, *Col6a3* levels were lower in ETPKO-sham kidneys compared to WT-sham kidneys, which may partially explain the reduction of *Col6a3* in ETPKO kidneys under IRI conditions. In contrast, on day 12, *Col6a3* expression was comparable in sham kidneys from WT and ETPKO mice but significantly reduced in ETPKO-IRI kidneys compared to WT-IRI kidneys (Fig. 3d). These findings suggest that *Etp* depletion diminishes *Col6a3* expression following IRI and that ETP may act as an upstream positive feedback signal for *Col6a3*. To evaluate additional fibrotic markers, RT-qPCR was performed for a panel of genes at days 7 and 12 post-IRI. No significant changes were observed at day 7 (Fig. 3e), whereas most fibrotic genes were significantly downregulated at day 12 in ETPKO-IRI kidneys compared to WT-IRI kidneys (Fig. 3f). In line with these transcriptional changes, fibrotic protein expression was more effectively downregulated in ETPKO-IRI kidneys at day 12 compared to day 7 (Fig. 3g). For example, on day 7, α-SMA expression was elevated in ETPKO-IRI kidneys, possibly as a compensatory response to ETP depletion. However, by day 12, α-SMA expression was significantly reduced in ETPKO-IRI kidneys compared to WT-IRI kidneys (Fig. 3g). COL6 was highly upregulated in both WT-IRI and ETPKO-IRI kidneys on day 7, but its expression was minimal or absent in ETPKO-IRI kidneys by day 12. Notably, mCherry expression was high at day 7 but significantly decreased by day 12 (Fig. 3g), indicating that ETP is highly upregulated at day 7 by IRI but its induction is substantially reduced by day 12 (Fig. 3g). These results suggest a sequential mechanism in which ETP upregulation in the early post-IRI phase drives fibrotic gene and protein expression, which becomes more pronounced by day 12, indicating a progressive fibrotic response. Collectively, these findings indicate that ETP depletion attenuates fibrotic gene expression in the kidney following IRI in a time-dependent manner.

**Fig. 3.**
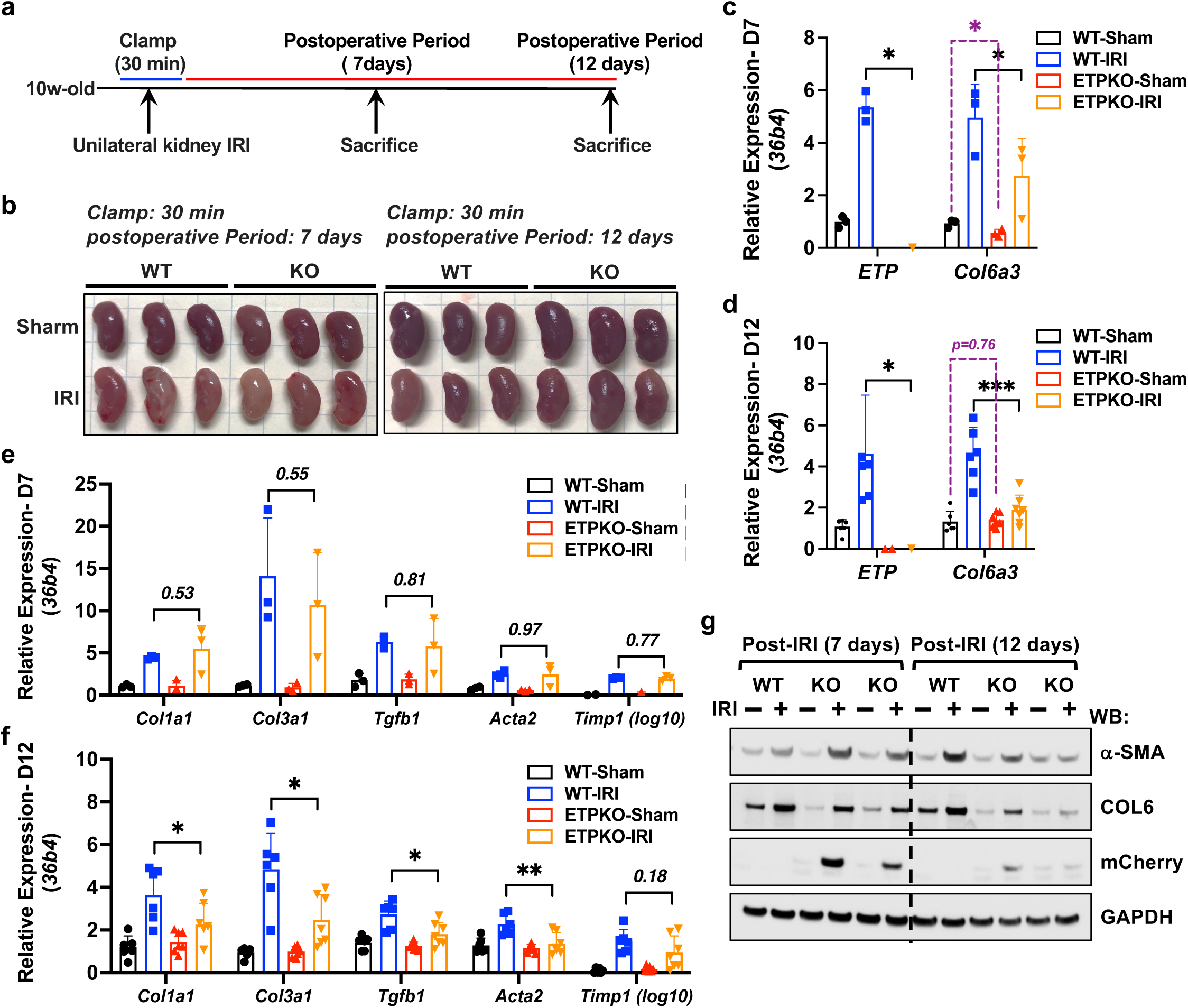
ETP depletion reduces local fibrotic gene mRNA expression following kidney ischemia-reperfusion injury. **a** Schematic representation of the experimental design for the unilateral kidney IRI model. The procedure involves a 30-minute clamp of the renal artery, followed by postoperative period of 7 or 12 days. **b** Representative images of sham and post-IRI kidneys at specified time points (n = 3 male mice/group). **c-d** *Col6a3* and *Etp* mRNA expression at day 7 (c) and day 12 (d) post-IRI, normalized to *36b4*. Data are presented as the mean± SEM (n = 3-6 male mice/group) and was analyzed by two-tailed Student’s t-tests. *, p<0.05; ** p<0.01. **e-f** Fibrotic gene mRNA expression at day 7 (e) and day 12 (f) post-IRI, normalized to *36b4*. Data are presented as the mean± SEM (n = 3-6 male mice/group) and was analyzed by two-tailed Student’s t-tests. *, p<0.05; ** p<0.01. **g** Immunoblot analysis of fibrotic protein levels in kidney tissues from WT and ETPKO mice under sham and post-IRI conditions (n = 3 male mice/group)

### Reduced expression of fibrotic proteins in ETP KO mice in unilateral kidney ischemia-reperfusion injury models

As we observed high ETP levels on day 7 and lower fibrotic protein levels on day 12, further analyses of fibrotic protein expression were conducted on day 12 post-IRI. Body weights and kidney weights were measured on day 12, and no significant changes were observed between sham and post-IRI mice (Fig. 4a, b). Analysis of fibrotic protein levels, including COL1, α-SMA, and COL6, revealed that these proteins were highly upregulated in WT-IRI kidneys compared to WT-sham kidneys (Fig. 4c). In contrast, COL1 and α-SMA were only slightly upregulated in ETPKO-IRI kidneys compared to ETPKO-sham kidneys, whereas COL6 expression levels remained unchanged between ETPKO-IRI and ETPKO-sham kidneys (Fig. 4c). Importantly, these fibrotic proteins were substantially reduced in ETPKO-IRI kidneys compared to WT-IRI kidneys, indicating that ETP promotes the induction of fibrotic proteins under IRI conditions. We have previously shown that mCherry (reflecting ETP) expression was substantially downregulated on day 12 compared to day 7 post-IRI (Fig. 3g). Following overexposure to detect the mCherry signal on day 12 post-IRI, mCherry was indeed still detectable in ETPKO mice (Fig. 4c). Among the fibrotic markers, COL6 showed a distinct pattern of regulation. Its expression was lower in sham kidneys from ETPKO mice compared to WT mice, suggesting that ETP depletion reduces COL6 expression independently of IRI. However, COL6 was highly upregulated in WT-IRI kidneys but not in ETPKO-IRI kidneys, emphasizing the critical role of ETP as a regulator of COL6 protein expression as a function of IRI (Fig. 4c). Subsequently, we performed IF analysis of mCherry, ETP, α-SMA, and COL6 in the kidney to corroborate our immunoblot analyses (Fig. 4d and Supplementary Fig. 4a-f). As expected, mCherry expression was only weakly detectable, and ETP was barely detectable in the IRI kidneys on day 12 (data not shown). Therefore, we used day 7 kidneys for IF analysis of mCherry and ETP. We observed increased mCherry staining in ETP-IRI kidneys, reflecting ETP upregulation on day 7 following IRI. We noted background signals from the ETP antibody that we generated in all kidney samples (highlighted with white arrows), including those from ETPKO mice. However, we observed a specific ETP signal (highlighted with yellow arrows) only in WT-IRI kidneys (Fig. 4d; showing both the background signal and the ETP signal together, and Supplementary Fig. 4e; highlighting only ETP signal-enriched regions). Additionally, we assessed ETP expression in various tissues under unilateral kidney IRI models. ETP was detected exclusively in subcutaneous and epididymal white adipose tissue (sWAT and eWAT) from WT mice but not in ETPKO mice (Supplementary Fig. 5a, b). While ETP is known to be highly expressed under high-fat diet conditions,^22^ the kidney IRI model was conducted under normal chow diet conditions, resulting in only a limited amount of ETP signal detectable in both fat tissues. In other tissues, such as the liver and lung, ETP signals were undetectable in both WT and ETPKO mice, with only non-specific signals observed (Supplementary Fig. 5a). Consistent with our immunoblotting results (Fig. 4c), α-SMA and COL6 expression were strongly upregulated on day 12 in WT-IRI kidneys compared to WT-sham kidneys (Fig. 4d). Notably, IRI in WT mice induced robust COL6 deposition in the ECM, a hallmark of fibrosis. In contrast, COL6 deposition was completely abolished in the kidneys of ETPKO mice under IRI conditions, indicating that ETP is a critical regulator of fibrosis through its role in collagen deposition. These findings further highlight the essential role that ETP plays in maintaining COL6 expression and driving fibrotic remodeling. Although α-SMA and COL6 expression were also upregulated on day 12 in ETPKO-IRI kidneys compared to ETPKO-sham kidneys, their upregulation was diminished compared to what is seen in WT-IRI kidneys (Fig. 4e, f). Overall, these findings demonstrate that ETP depletion significantly reduces fibrotic protein expression, particularly on day 12 post-IRI, highlighting its crucial role in driving kidney fibrosis following injury.

**Fig. 4.**
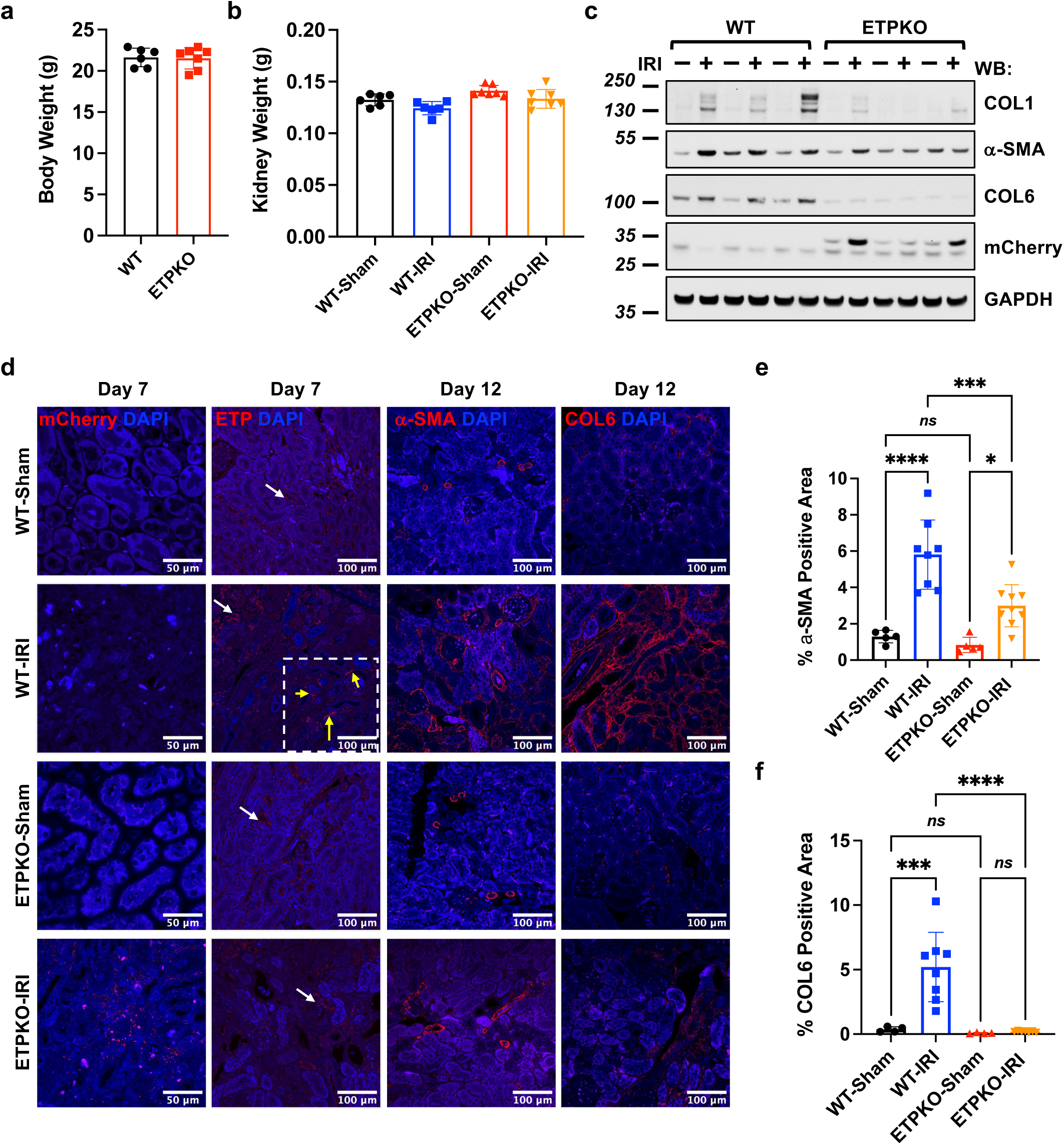
ETP depletion reduces local fibrotic gene protein expression following kidney ischemia-reperfusion injury. a-b. Body weight (a) and kidney weight (b) of mice at day 12 post-IRI (n = 6-7 male mice/group). **c** Immunoblot analysis of fibrotic protein levels in kidney tissues from WT and ETPKO kidney mice under sham and post-IRI conditions (n = 3 male mice/group). **d** Representative IF images of mCherry, ETP, α-SMA, and COL6 across kidney tissues from WT and ETPKO mice under sham and post-IRI conditions (n = 3 male mice/group). For the ETP staining, instances of non-specific signal (white arrows) and specific signal (yellow arrows) are indicated. **e-f** Quantification of α-SMA (e) and COL6 (f) positive areas (%), based on the staining shown in (d). Data are presented as the mean± SEM and was analyzed by one-way ANOVA. *ns*, non-significant; *, p<0.05; *** p<0.001; **** p<0.0001.

### ETP Depletion Prevents Overt Local Fibrosis Following Kidney Ischemia-Reperfusion Injury

Beyond fibrotic protein levels, we assessed tissue fibrosis using established histological methods. As revealed by picrosirius red staining of entire kidney sections, tissue fibrosis was significantly induced in WT kidneys following IRI (Fig. 5a). In contrast, kidneys from ETPKO mice subjected to IRI exhibited a pronounced reduction in fibrosis (Fig. 5a). Only minimal fibrosis was observed in WT-sham and ETPKO-sham groups. Quantification of picrosirius red-positive areas (%), based on the staining results (Fig. 5a), demonstrated that fibrosis was strongly induced in WT-IRI kidneys but significantly reduced in ETPKO-IRI kidneys (Fig. 5b). Of note, physical damage or membrane areas of the kidney resulted in some artificial positive signals, marked with black asterisks, which were excluded from quantification. Representative images of picrosirius red-stained kidney tissues from all experimental groups supported our observations (Fig. 5c). Collectively, these results demonstrate that the absence of ETP correlates with reduced tissue fibrosis in the kidney following IRI, suggesting that ETP is a critical factor in this process.

**Fig. 5.**
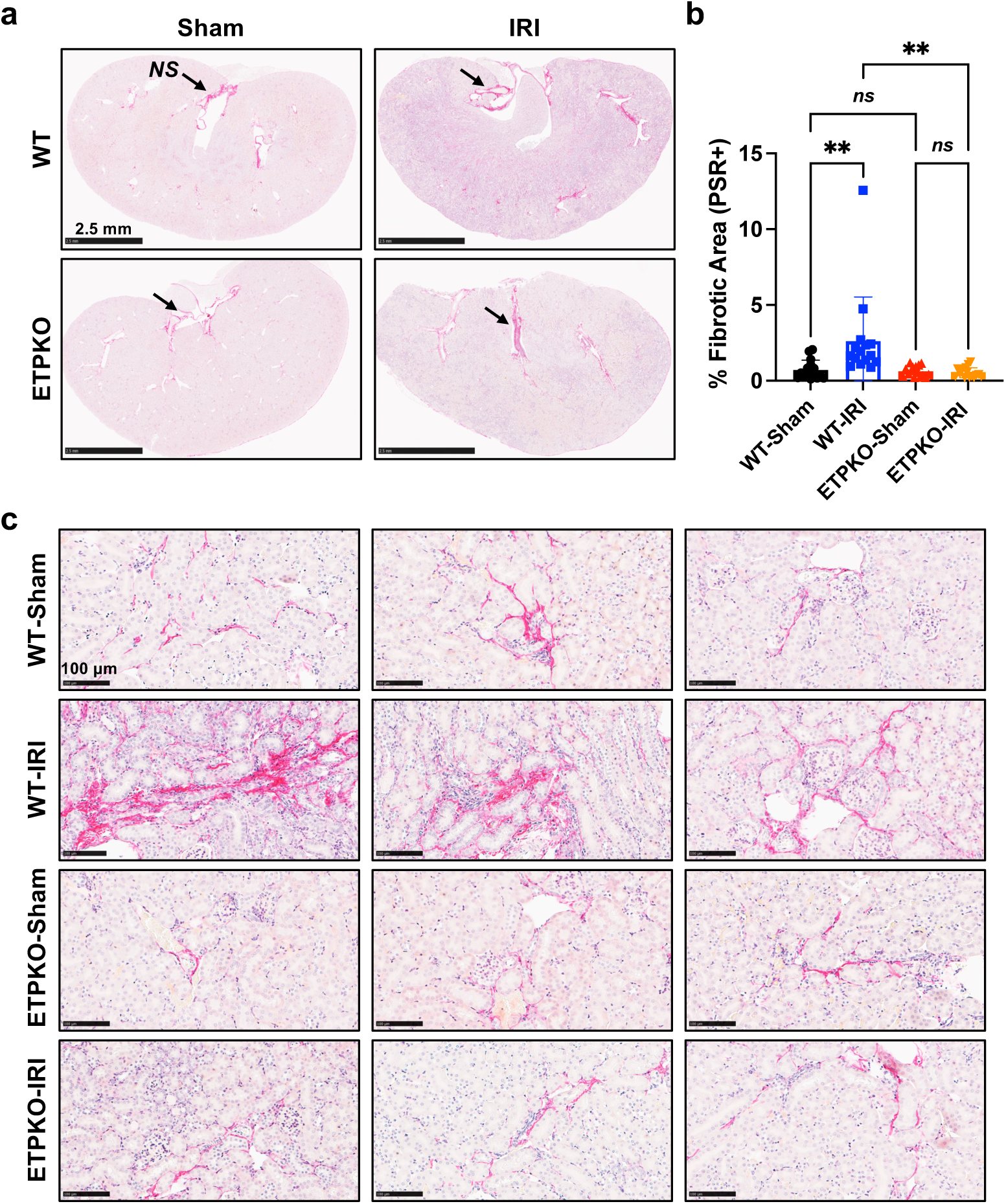
ETP depletion prevents overt local fibrosis following kidney ischemia-reperfusion injury. **a** Picrosirius red staining showing fibrosis in the entire kidney across experimental groups (n = 3 male mice/group). Areas of n**on-specific staining (black arrows) were** excluded from quantification. Scale bar equals 2.5 mm. **b** Quantification of picrosirius red-positive areas (%), based on the staining results shown in (a). Data are presented as the mean± SEM and was analyzed by one-way ANOVA. ***ns*, non-significant;** ** p<0.01. **c** Representative images of picrosirius red-stained kidney tissues from all experimental groups. Scale bar equals 100 μm.

### ETP Ablation Alleviates Proteinuria in a Two-Stage Bilateral Kidney Ischemia-Reperfusion Injury Model

In our previous study using ETP-neutralizing antibodies in the POD-ATTAC model, neutralizing ETP significantly ameliorated renal fibrosis and improved renal function, as reflected by reduced proteinuria ^15^. To investigate whether the genetic ablation of ETP similarly protects mice from severe kidney dysfunction following injury, we performed a two-stage bilateral kidney IRI ^20^. We first subjected the left kidney to IRI, followed by a recovery period of 7 days. Subsequently, we performed a second round of IRI on the right kidney and monitored postoperative outcomes for an additional seven days (Fig. 6a). Urine samples were collected at the indicated time points to measure proteinuria levels. On day 14, we observed that the right kidney became larger than the left kidney in WT-IRI mice (Fig. 6b). However, in ETPKO mice, kidney sizes remained comparable (Fig. 6b). Importantly, on day 10, albumin levels in urine samples were significantly reduced in ETPKO mice compared to WT mice, indicating that ETPKO mice exhibited reduced proteinuria and recovered faster (Fig. 6c). These findings suggest a protective role of ETP depletion in preventing renal dysfunction. Histological analysis of the first IRI-injured left kidney revealed significant fibrosis in WT mice, whereas ETPKO mice exhibited markedly reduced fibrosis, consistent with previous results (Fig. 6d, e). However, no significant fibrosis differences were observed in the second IRI-injured right kidney (Fig. 6d, e). In our previous experiments (Figure 3), post-operative samples collected on day 7 did not show significant differences in fibrotic gene expression, but by day 12, significant differences were observed. In our two-stage bilateral IRI model, fibrosis in the first IRI kidney was allowed to progress to day 14, whereas fibrosis in the second IRI kidney was allowed to progress only to day 7. This difference in progression time likely contributed to the observed variations in fibrosis outcomes between the left (first IRI) and right (second IRI) kidneys of the mice. Based on these and previous findings, we confirmed that days 12–14 post-IRI represent an optimal time period for detecting fibrosis differences, whereas day 7 is not as informative in this model. These results demonstrate that ETP depletion exerts a protective effect on renal function in the kidney IRI model by ameliorating the fibrotic response.

**Fig. 6.**
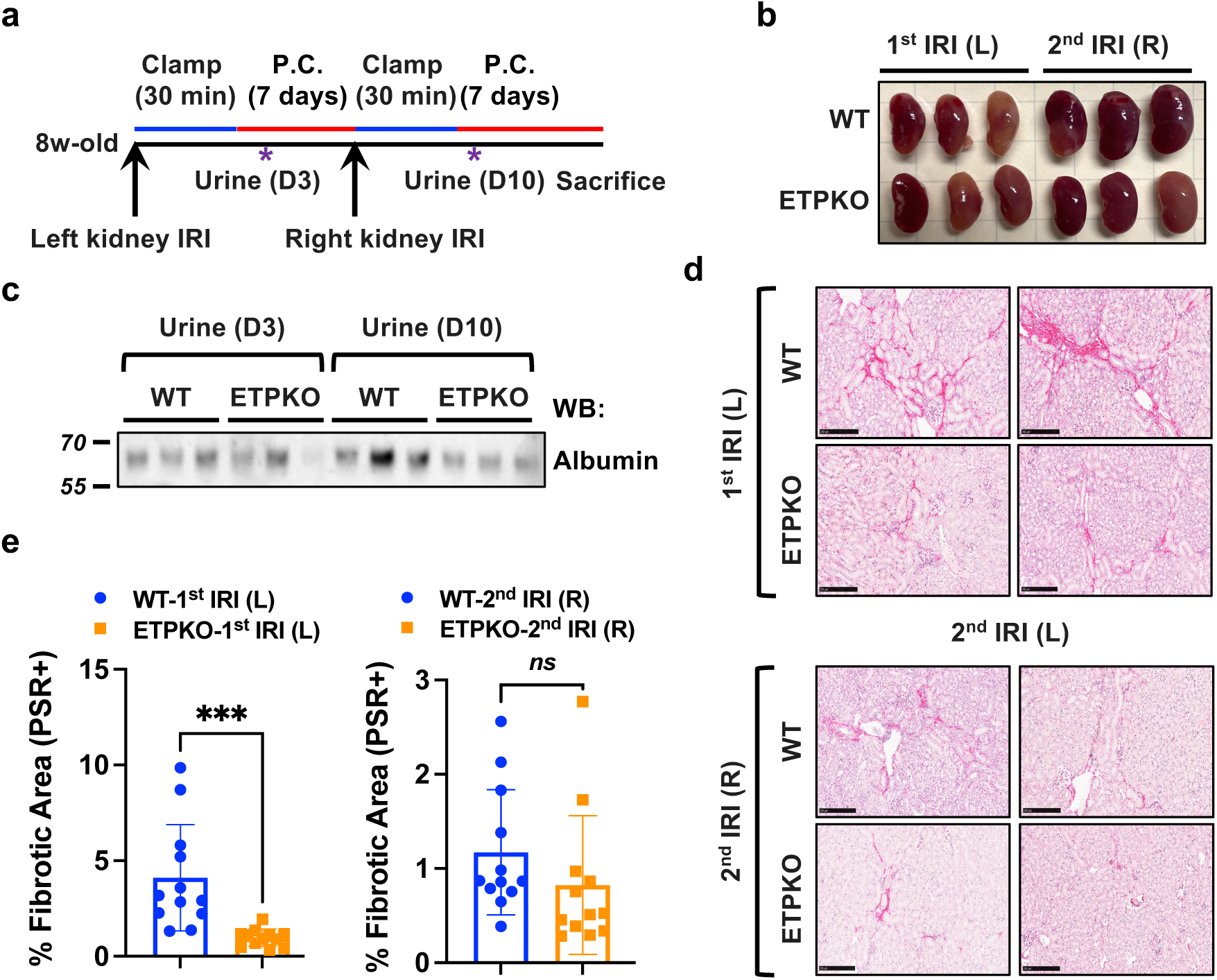
ETP KO mice exhibited a reduced proteinuria phenotype in a two-stage bilateral ischemia-reperfusion injury model. **a** Schematic representation of the experimental setup for the two-stage bilateral kidney ischemia-reperfusion injury (IRI) model. The procedure involves a 30-minute clamping of the left kidney’s renal artery, followed by post-operative period of 7 days. Subsequently, a 30-minute clamping of the right kidney’s renal artery is performed, followed by a post-operative period of another 7 days. Urine samples were collected at the indicated time points. **b** Representative images post-IRI kidneys at day 14 (n = 3 male mice/group). **c** Immunoblot analysis of albumin levels in urine samples from WT and ETPKO mice (n = 3 male mice/group). **d** Picrosirius red staining showing fibrosis in the kidney across experimental groups (n = 3 male mice/group). Scale bar equals 250 μm. **e** Quantification of picrosirius red-positive areas (%), based on the staining shown in (d). Data are presented as the mean± SEM and were analyzed by two-tailed Student’s t-tests. *ns*, non-significant; *** p<0.001.

## DISCUSSION

Fibrosis is a pathological process characterized by the excessive accumulation of ECM components in organs such as the heart, liver, and kidneys, often resulting from chronic injury or inflammation ^23–25^. It is a critical feature of various disease models, including chronic kidney disease ^26^, liver fibrosis ^27^, and cardiac fibrosis ^28^. ETP has been implicated in fibrotic processes across various tissues, including the heart, liver, and kidneys, highlighting its overall significance in pathogenesis ^1,12,20^. Elevated ETP levels are strongly associated with fibro-inflammatory diseases such as chronic kidney disease ^29^, liver disease ^21^, cardiovascular disease, and heart failure ^30^, disease states that are frequently seen in combination and referred to as the cardiorenal metabolic syndrome. In this study, we explored the critical role of ETP in kidney fibrosis by generating whole-body ETPKO mice. Unlike traditional *Col6a3* mutant models, whose phenotypes appear dominated by the present disruption of COL6A3 expression and function ^8^ ^9^, our ETPKO model specifically eliminates ETP expression with only modest effects on COL6A3 expression. This specificity allowed us to investigate the role of ETP in fibrosis without having to account for profound confounding effects arising from the impairment of COL6A3. Utilizing a unilateral kidney IRI model in ETPKO mice, we demonstrated that ETP ablation significantly reduces both fibrotic gene expression and overall tissue fibrosis in the kidneys following injury. These findings align with our previous studies using an ETP-neutralizing antibody in the POD-ATTAC model, where ETP neutralization significantly ameliorated renal fibrosis and improved renal function, as indicated by reduced proteinuria^15^. Consistent with these observations, in a two-stage bilateral kidney IRI model, we further demonstrate that genetic ETP depletion prevents injury-induced deterioration of renal function by mitigating fibrosis. Our antibody-mediated ETP inhibition ^15^, as well as genetic ETP depletion studies, both provide strong evidence that ETP is a critical regulator of kidney fibrosis and underscore the potential of ETP as a therapeutic target in fibrotic diseases. Interestingly, our findings reveal a significant downregulation of COL6 protein in ETPKO mice compared to WT mice, regardless of IRI, potentially arising as a secondary effect of ETP depletion. Further research is needed to elucidate the molecular mechanisms by which ETP influences COL6 expression. Our study herein highlights the utility of the ETPKO mouse model as a robust tool for investigating the physiological and pathological roles of ETP. For the first time, we provide experimental evidence from a genetic mouse model that ETP plays a crucial role in the fibrotic process, establishing our new ETPKO model as a valuable resource for studying the involvement of ETP in other pathological conditions, such as metabolic disorders, fibro-inflammatory diseases, cardiac diseases, and cancer. Additionally, through mCherry-based immunoblotting and immunostaining, we suggest that this approach could be utilized to track COL6A3 and thus ETP expression across different cell types and tissues under physiological and pathological conditions. This approach provides a valuable opportunity to better understand the patterns of ETP expression over time and across different tissues, offering the foundation for future investigations into the role of ETP in various disease models. Taking full advantage of the original Cre-dependent *Col6a3-ETP+mCherry-CAAX* allele, future studies should focus on generating tissue-specific ETPKO mice to address the contributions of different cell types to overall ETP levels. This will help elucidate how ETP expression changes and how it impacts different outcomes in various tissues. Collectively, our results highlight ETP as a critical regulator of fibrosis and suggest its potential as both a biomarker for tracking fibrotic progression and a therapeutic target for the treatment of fibrotic diseases.

## ACKNOWLEDGEMENTS

We thank the UTSW Transgenic Core, and Animal Resource Center, and ARC Diagnostic Lab. We thank the UTSW Quantitative Light Microscopy Core Facility and the UTSW Whole Brain Microscopy Facility (RRID:SCR_017949) for providing imaging equipment. Katherine Luby-Phelps at the UTSW Quantitative Light Microscopy Core Facility was supported by NIH grant 1S10OD021684-01. This work was supported by NIH grants RC2-DK118620, R01-DK55758, R01-DK099110, R01-DK127274 and R01-DK131537 to P.E.S

## DATA AVAILABILITY

The datasets and unique research materials generated during the current study are available from the corresponding author on reasonable request.

