## Supplemental Information for "ETP-Specific Knockout Mice Reveal Endotrophin as a Key Regulator of Kidney Fibrosis in Ischemia-Reperfusion Injury Models"

Conflict of interest: The authors have declared that no conflict of interest exists.

***List of Supplement Materials***

1. Supplementary Figures S1 to S5

2. Tables S1 to S4

3. Supplemental Information

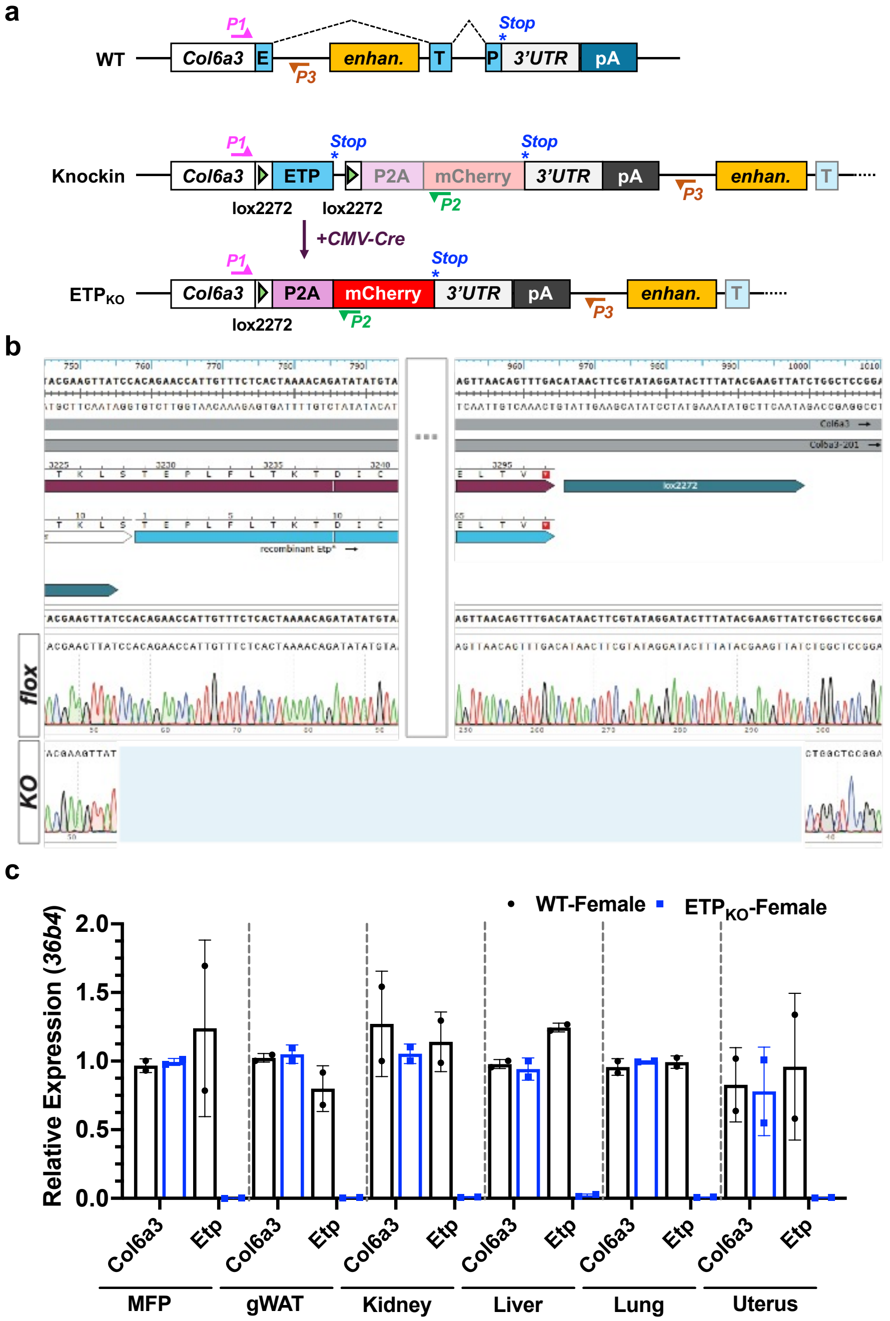


**Supplementary Fig. 1 Generation and validation of whole body ETP knockout mice.**
**a** Schematic representation of the wild-type Col6a3 allele and the new Col6a3-ETP+mCherry-CAAX allele before and after Cre-mediated recombination. The stop codons of the ETP and P2A-mCherry-CAAX reading frames are shown as asterisks (blue). **b** Sanger sequencing of genomic DNA from knock-in and KO mice to validate the successful deletion of the ETP-encoding DNA sequence. **c** Col6a3 and Etp mRNA expression, normalized to 36b4. Data are presented as the mean ± SEM (n = 2 female mice per group) and were analyzed by two-tailed Student's t-tests. *p < 0.05, **p < 0.01.

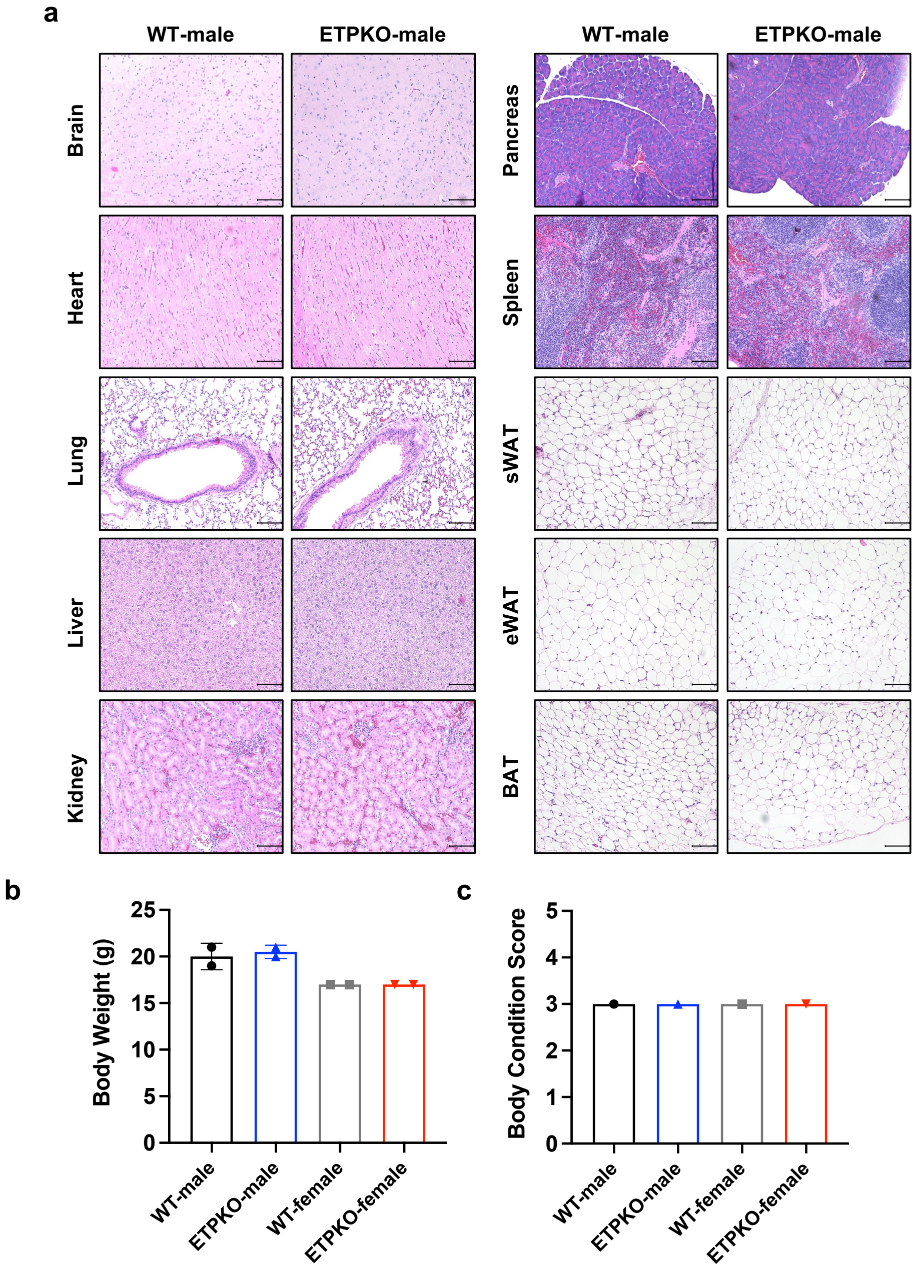


**Supplementary Fig. 2 Whole body ETP knockout mice display no overt phenotype in histopathological analysis.**

**a** Histopathological analysis of H&E-stained tissues from WT and ETPKO mice revealed normal findings (n = 2 male and 2 female mice per group). **b, c** Body weights (b) and body condition scores (c). Data are presented as the mean ± SEM (n = 2 male and 2 female mice per group).

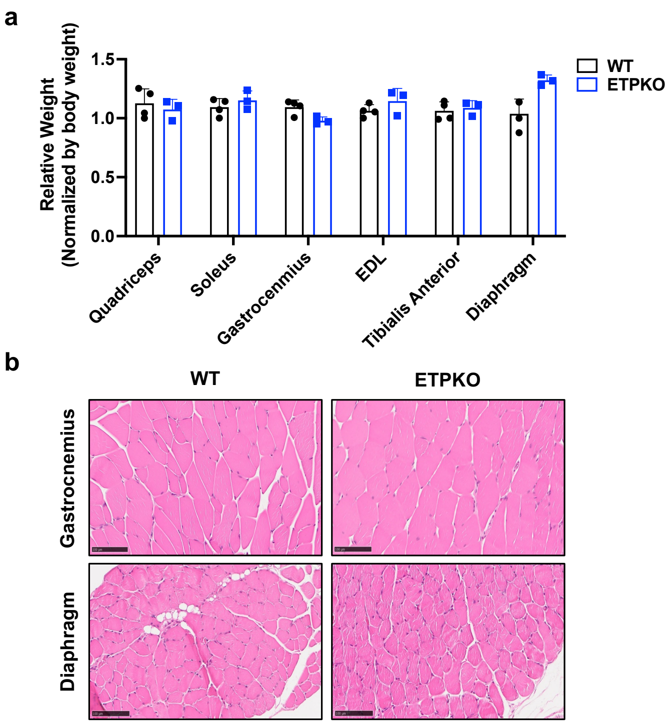


**Supplementary Fig. 3 Whole body ETP knockout mice display no signs of skeletal muscle defects.**

**a** Muscle weights of WT and ETPKO mice. Data are presented as the mean ± SEM (n = 3-4 male mice per group; 10 weeks old). **b** Histological analysis of H&E-stained gastrocnemius and diaphragm muscles in WT and ETPKO mice revealed normal findings (n = 3-4 mice per group; 10 weeks old).

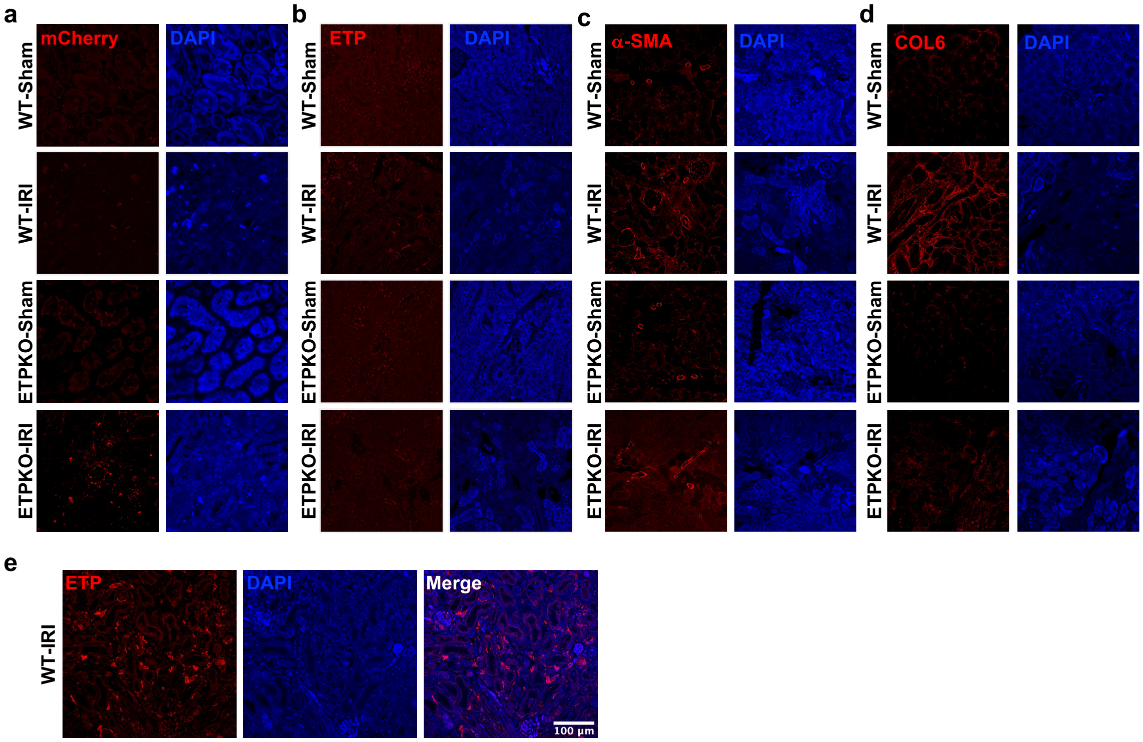


**Supplementary Fig. 4 ETP depletion reduces local fibrotic gene protein expression following kidney ischemia-reperfusion injury.**
**a-d** Representative IF images of mCherry (a), ETP (b), α-SMA (c), and COL6 (d) across kidney tissues from WT and ETPKO mice under sham and post-IRI conditions (n = 3 male mice per group). **e** Additional IF images showing ETP expression in post-IRI kidneys of WT mice.

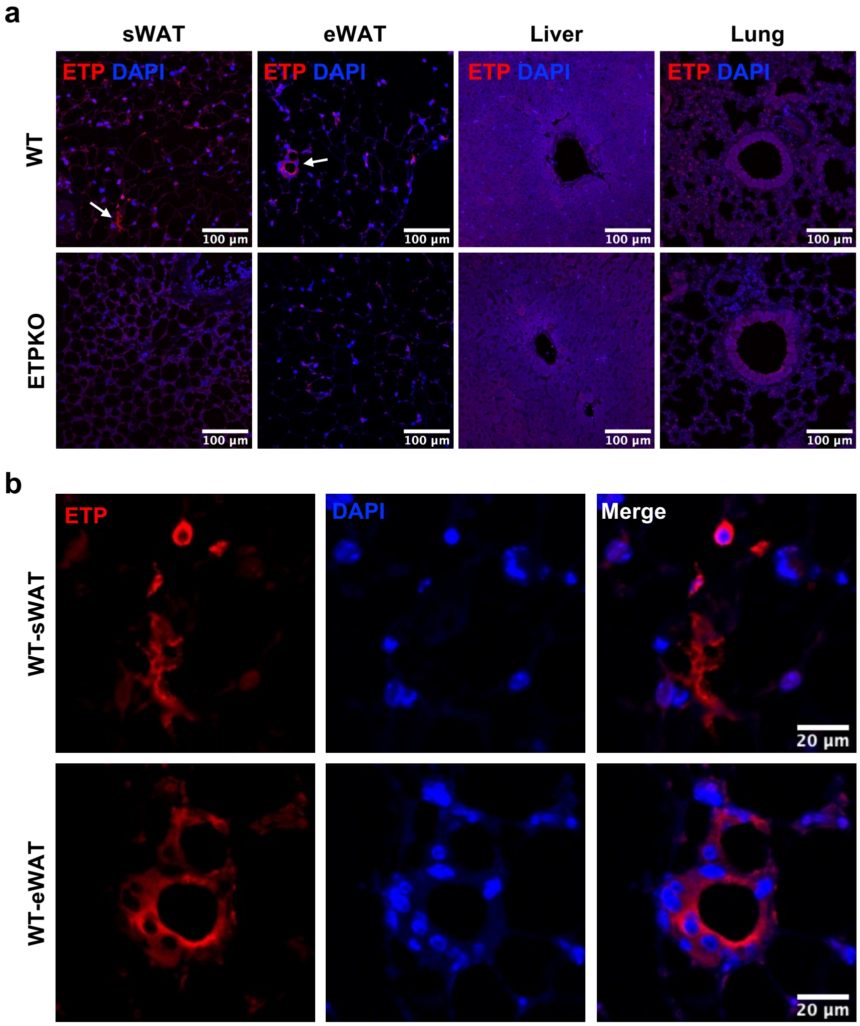


**Supplementary Fig. 5 ETP expression in multiple tissues following unilateral kidney ischemia-reperfusion injury.**

**a** Representative IF images of ETP across subcutaneous white adipose tissue (sWAT), epididymal white adipose tissue (eWAT), liver, and lung from WT and ETPKO mice post-IRI. Areas of magnification are indicated. **b** Magnified IF staining (white arrows; displayed in panel a) highlighting ETP expression in the sWAT and eWAT of WT mice post-IRI.

### Table 1. WT and ETP KO mice display normal phenotypes in histopathological analysis

| Animal ID | Gross Findings | Organ | Microscopic Findings |
|---|---|---|---|
| All WT and KO (male and female) | No lesions found | Brain | Normal |
| All WT and KO (male and female) | No lesions found | Heart | Normal |
| All WT and KO (male and female) | No lesions found | Lungs | Normal |
| All WT and KO (male and female) | No lesions found | Liver | Normal |
| All WT and KO (male and female) | No lesions found | Kidneys | Normal |
| All WT and KO (male and female) | No lesions found | Spleen | Normal |
| All WT and KO (male and female) | No lesions found | Pancreas | Normal |
| All WT and KO (male and female) | No lesions found | sWAT | Normal |
| All WT and KO (male and female) | No lesions found | gWAT | Normal |
| All WT and KO (male and female) | No lesions found | brown AT | Normal |

**Table 2. List of genotyping primer sequences**

| Name | Forward primer | Reverse primer |
|---|---|---|
| P1 (G336) | TCTTCAGGCAGCACACCGAG | |
| P2 (G337) | | TCACCATAGGACCGGGGTTTT |
| P3 (G338) | | CTGAGGACCCCTTTGGAACTG |
| CMV-Cre | GCGGTCTGGCAGTAAAAACTATC | GTGAAACAGCATTGCTGTCACTT |

### Table 3. List of primers used for qPCR

| Gene name | Forward primer | Reverse primer |
|---|---|---|
| *Etp* | GAAAGGGGATTATGGCTCAGG | TCACTGTCCCAGCATCTTGTG |
| *Col6a3* | GCAACTGTTCCTGAACTCAACT | ATCTTTTGGGGTCCGTCAACT |
| *Col1a1* | GTGCTCCTGGTATTGCTGGT | GGCTCCTCGTTTTCCTTCTT |
| *Col3a1* | GGGTTTCCCTGGTCCTAAAG | CCTGGTTTCCCATTTTCTCC |
| *Tgfb1* | ACCATGCCAACTTCTGTCTG | CGGGTTGTGTTGGTTGTAGA |
| *Acta2* | GTACCACCATGTACCCAGGC | GCTGGAAGGTAGACAGCGAA |
| *Timp1* | CCCCAGAAATCAACGAGACCA | ACTCTTCACTGCGGTTCTGG |
| *36B4* | AGATTCGGGATATGCTGTTGGC | TCGGGTCCTAGACCAGTGTTC |

### Table S4. Resources table

| REAGENT or RESOURCE | SOURCE | IDENTIFIER |
|---|---|---|
| **ANTIBODIES** | | |
| ETP | Home-made | Cat #9661 |
| a-SMA | Cell Signaling | Cat #19245 |
| COL6 | Invitrogen | Cat #MA5-32412 |
| COL1 | SouthernBiotech | Cat # 1310-01 |
| mCherry & RFP | Cell Signaling & Rockland | Cat #43590 & 600-401-379 |
| Alexa Fluor 488 | Invitrogen | Cat #A-11006 |
| Alexa Fluor 594 | Invitrogen | Cat #A-11037 |
| **CHEMICALS and OTHERS** | | |
| PowerUp™ SYBR™ Green Master | Applied Biosystems | Cat #A25742 |
| Normal goat serum | ThermoFisher | Cat #31873 |
| RIPA Buffer | Pierce | Cat #89900 |
| 4-12% gradient polyacrylamide-SDS gel | Invitrogen | Cat #NP0336 |
| Nitrocellulose membrane | BioRad | Cat #1704159 |
| PrimeScript™ RT Master Mix | TaKaRa | Cat #RR036A |
| Antigen Unmasking Solution, Citrate-Based | Vector Labs | Cat #H-3300-250 |
| VECTASHIELD mounting medium with DAPI | Vector Labs | Cat #H-2000 |
| BSA | Sigma-Aldrich | Cat #A3294 |
| GoTaq G2 Green Master Mix | Promega | Cat #M7823 |
| TRIzol™ Reagent | Invitrogen | Cat #15596026 |
| SuperSignal West Pico PLUS Chemiluminescent Substrate | ThermoFisher | Cat #34577 |
| Protease Inhibitor Cocktail | Millipore sigma | Cat #11873580001 |
| Phosphatase inhibitor cocktail 3 | Sigma | Cat #P0044 |
| Wiegert's Hematoxylin Solution | Sigma | Cat #HT1079 |
| **Kits** | | |
| RNA purification kit | Qiagen | Cat #74104 |
| EZ-10 DNAaway RNA miniprep kit | BIO BASIC | Cat #BS88136 |
| Picrosirius Red Stain Kit | Polyscience | Cat #24901 |
| H&E Staining Kit | Abcam | Cat #ab245880 |
| **EXPERIMENTAL MODELS: ORGANISMS/STRAINS** | | |
| *ET KO* mice | This study | N/A |
| *CMV-Cre* mice | Teppei Fujikawa provided | N/A |
| **OLIGONUCLEOTIDES** | | |
| Primers, see Table S1, and S2 | | |
| **Software and Algorithms** | | |
| FIJI/ImageJ | NIH | RRID: SCR_003070 |
| Prism10 | GraphPad | RRID:SCR_002798 |

### Supplemental Information

*pUC57-Col6a3-Etp+mCherry-CAAX*

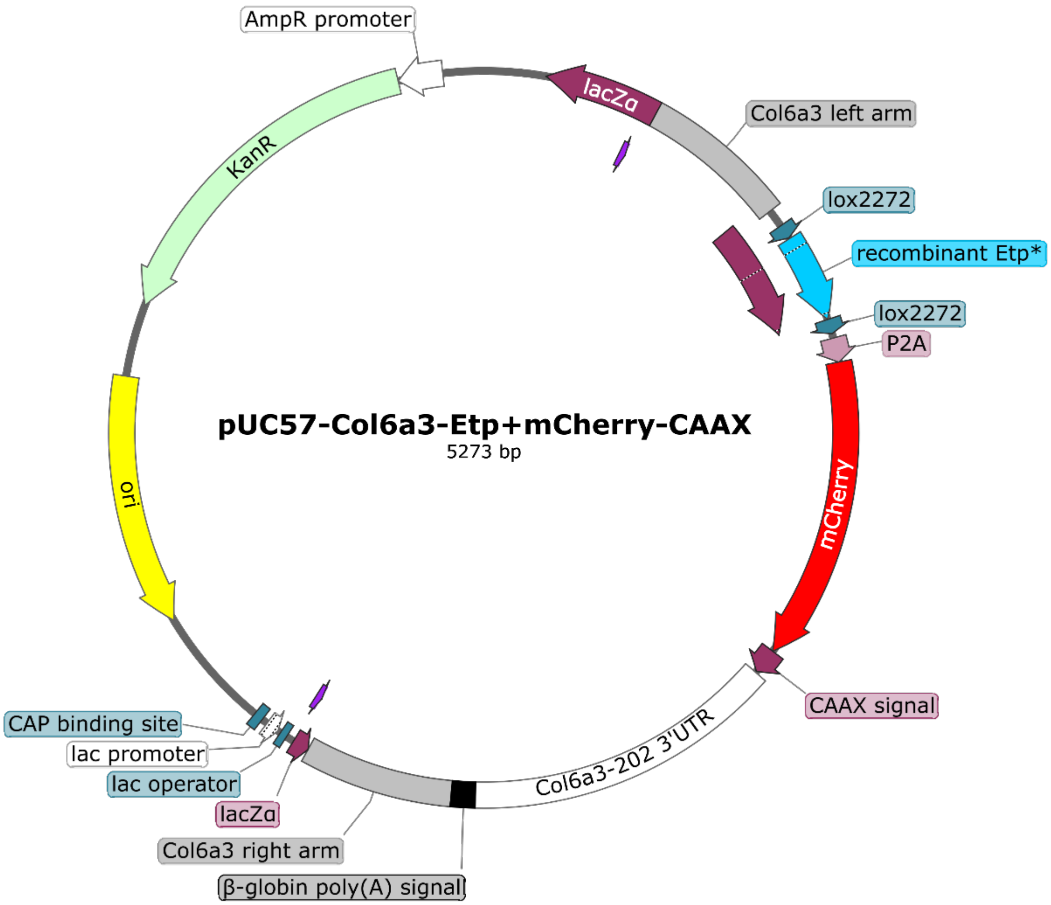


TCGCGCGTTTCGGTGATGACGGTGAAAACCTCTGACACATGCAGCTCCCGGAGACTGTCA
CAGCTTGTCTGTAAGCGGATGCCGGGAGCAGACAAGCCCGTCAGGGCGCGTCAGCGGGT
GTTGGCGGGTGTCGGGGCTGGCTTAACTATGCGGCATCAGAGCAGATTGTACTGAGAGTG
CACCATATGCGGTGTGAAATACCGCACAGATGCGTAAGGAGAAAATACCGCATCAGGCGC
CATTCGCCATTCAGGCTGCGCAACTGTTGGGAAGGGCGATCGGTGCGGGCCTCTTCGCTA
TTACGCCAGCTGGCGAAAGGGGGATGTGCTGCAAGGCGATTAAGTTGGGTAACGCCAGG
GTTTTCCCAGTCACGACGTTGTAAAACGACGGCCAGTGAATTGACGCGTATTGGGATAAAC
AATGATGCCTTGAGTCCATTCTGAAGCACAGACACAATTAAGTACAATCCATCAATCTTGGG
TACTTAGGCTACACCCTGAAAATCAAAATCACAGCAACAGCATAGGAGACAGGGTTCCCTG
TGCCATATGCTCTCATTCTTGGTTCCTGGGACCCTTTAAGGGGGAGACCAGATTTGCTCTG
CCTTAGCCTATCATGTCTAGGTGGCATGTGGGAAATGGGGACCTGGATCCTGGCGAGGAA
GATAGTAGGATCTCTGTTGCCACTTTTAGTTTCCGAGTCAACTACTAGTTGGTTGTATTTTC
TTTTCTTATAGAGAAAACCCAGCCTCCACCTCTTCAGGCAGCACACCGAGCAAGCTCAAGT
ACAATCAATCTGATGGTGAACATAACTTCGTATAGGATACTTTATACGAAGTTATCCACAGA
ACCATTGTTTCTCACTAAAACAGATATATGTAAGCTGTCCAGAGATGCTGGGACTTGTGTG

GACTTCAAGTTACTATGGCACTATGACCTAGAGAGCAAAAGTTGCAAGAGATTCTGGTATG
GAGGTTGTGGAGGCAACGAGAACAGATTCCACTCCCAGGAAGAATGTGAAAAGATGTGTA
GTCCTGAGTTAACAGTTTGACATAACTTCGTATAGGATACTTTATACGAAGTTATCTGGCTC
CGGAGCCACGAACTTCTCTCTGTTAAAGCAAGCAGGAGACGTGGAAGAAAACCCCGGTCC
TATGGTGAGCAAGGGCGAGGAGGATAACATGGCCATCATCAAGGAGTTCATGCGCTTCAA
GGTGCACATGGAGGGCTCCGTGAACGGCCACGAGTTCGAGATCGAGGGCGAGGGCGAG
GGCCGCCCCTACGAGGGCACCCAGACCGCCAAGCTGAAGGTGACCAAGGGTGGCCCCCT
GCCCTTCGCCTGGGACATCCTGTCCCCTCAGTTCATGTACGGCTCCAAGGCCTACGTGAA
GCACCCCGCCGACATCCCCGACTACTTGAAGCTGTCCTTCCCCGAGGGCTTCAAGTGGGA
GCGCGTGATGAACTTCGAGGACGGCGGCGTGGTGACCGTGACCCAGGACTCCTCCCTGC
AGGACGGCGAGTTCATCTACAAGGTGAAGCTGCGCGGCACCAACTTCCCCTCCGACGGC
CCCGTAATGCAGAAGAAGACCATGGGCTGGGAGGCCTCCTCCGAGCGGATGTACCCCGA
GGACGGCGCCCTGAAGGGCGAGATCAAGCAGAGGCTGAAGCTGAAGGACGGCGGCCAC
TACGACGCTGAGGTCAAGACCACCTACAAGGCCAAGAAGCCCGTGCAGCTGCCCGGCGC
CTACAACGTCAACATCAAGTTGGACATCACCTCCCACAACGAGGACTACACCATCGTGGAA
CAGTACGAACGCGCCGAGGGCCGCCACTCCACCGGCGGCATGGACGAGCTGTACAAGAA
GCTGAACCCTCCTGATGAGAGTGGCCCCGGCTGCATGAGCTGCAAGTGTGTGCTCTCC**TA**
**A**CAAGAGCCTAAGCATGGCCTTCAGGCAACACGTACCTCTGGGAGAAGGAGGAGGCAGC
CATTTCTAACTCGTTTCTATAGAAGCCCTGGGTAGATGCCTCAGCACGGTGCCTTTTCATG
CTTTGATTGACACTCAACCTCGGGAGGAAACCCTCTGCACGTGACCTGTCAATATGGTGCT
AAATGTGTCTATGGACCCTGCTCTCCGTCTCCAGGCAGTTCTACCGTATACTTGGACCCTT
GGGTTATAGCTAGCCACTGCTGGTGTTTATGTGAACATTCCTATAAATTCAATTTCCCTCTG
GAGTTCCACGCTACGCCTGTGCCAGGCAAACCCTGTGCCTAGAACATAGCCTGGACGTCA
CAGCTACTCTGTACATTTCTGCTTGGTTCATTCCTCTGTAGTTGCACGGCTTAGATGGAGAA
ACAAGAGTCTAACCTTCTCATGGTCCCAGTTTTCTGGATTAGACTTCGATCAATATTCTTCT
AAATCCTCTGACAAATGATCTAATTAGAAGAAATCAGACCTCTTTCCTGTGTGCATTGCTGG
GACAAATGCCTCCATTAGAAAATTCAAAGAAAGTCATAATCGAGAATCTCTTTGGTGGTCCT
CTAAGGCGGGTTGTTTTTCAATGTTGTTGTCTTGGAGCTTGGAGGTGAAATTCAATGTTTAA
AATTTTTAGGAAATTTATACAAAGAAACTTTTAAATAAAGTATATTGAATGTGCCATGAAATA
AAGGAAATTTATTTTCATTGCAATAGTGTGTTGGAATTTTTTGTGTCTCTCAGTAAGTCGTGG
CAGTACTAGGTCCCCTAATGGACTTCAGAAAGCATTCTGAGACCAGGGAGAAGTACTGCTT
AATGGCTTGCAAACCCTTTGAAATTGAGGCCACCTGCCTGAGACTAAATCACATGACTTCC
ATGGGATCAGTTCCAAAGGGGTCCTCAGCTGCTGGGTTTCATCTCACTCTACATGGTCTTG
GTTTCAAAAGCAGAGGAACCAAGCTTGTCACCATAGTGTTAGAAGAAGCTGGCTGGAGCT
GCTTCTGAGCCCATGGCTATTATGCACGGCTCCACCAGTCACTTCCTCTGTGTTATGTTGA
AACTGGCTAGGGTTGACTTGTGGTCTGTAGTGCTGATCCCAATGGCGCGCCGAGCTTGGC
TCGAGCATGGTCATAGCTGTTTCCTGTGTGAAATTGTTATCCGCTCACAATTCCACACAACA
TACGAGCCGGAAGCATAAAGTGTAAAGCCTGGGGTGCCTAATGAGTGAGCTAACTCACAT
TAATTGCGTTGCGCTCACTGCCCGCTTTCCAGTCGGGAAACCTGTCGTGCCAGCTGCATTA
ATGAATCGGCCAACGCGCGGGGAGAGGCGGTTTGCGTATTGGGCGCTGTTCCGCTTCCT
CGCTCACTGACTCGCTGCGCTCGGTCGTTCGGCTGCGGCGAGCGGTATCAGCTCACTCAA
AGGCGGTAATACGGTTATCCACAGAATCAGGGGATAACGCAGGAAAGAACATGTGAGCAA
AAGGCCAGCAAAAGGCCAGGAACCGTAAAAAGGCCGCGTTGCTGGCGTTTTTCCATAGGC
TCCGCCCCCCTGACGAGCATCACAAAAATCGACGCTCAAGTCAGAGGTGGCGAAACCCGA
CAGGACTATAAAGATACCAGGCGTTTCCCCCTGGAAGCTCCCTCGTGCGCTCTCCTGTTCC
GACCCTGCCGCTTACCGGATACCTGTCCGCCTTTCTCCCTTCGGGAAGCGTGGCGCTTTC
TCATAGCTCACGCTGTAGGTATCTCAGTTCGGTGTAGGTCGTTCGCTCCAAGCTGGGCTGT
GTGCACGAACCCCCCGTTCAGCCCGACCGCTGCGCCTTATCCGGTAACTATCGTCTTGAG
TCCAACCCGGTAAGACACGACTTATCGCCACTGGCAGCAGCCACTGGTAACAGGATTAGC

AGAGCGAGGTATGTAGGCGGTGCTACAGAGTTCTTGAAGTGGTGGCCTAACTACGGCTAC
ACTAGAAGAACAGTATTTGGTATCTGCGCTCTGCTGAAGCCAGTTACCTTCGGAAAAAGAG
TTGGTAGCTCTTGATCCGGCAAACAAACCACCGCTGGTAGCGGTGGTTTTTTTGTTTGCAA
GCAGCAGATTACGCGCAGAAAAAAAGGATCTCAAGAAGATCCTTTGATCTTTTCTACGGGG
TCTGACGCTCAGTGGAACGAAAACTCACGTTAAGGGATTTTGGTCATGAGATTATCAAAAA
GGATCTTCACCTAGATCCTTTTAAATTAAAAATGAAGTTTTAAATCAATCTAAAGTATATATG
AGTAAACTTGGTCTGACAGTTAGAAAAACTCATCGAGCATCAAATGAAACTGCAATTTATTC
ATATCAGGATTATCAATACCATATTTTTGAAAAAGCCGTTTCTGTAATGAAGGAGAAAACTC
ACCGAGGCAGTTCCATAGGATGGCAAGATCCTGGTATCGGTCTGCGATTCCGACTCGTCC
AACATCAATACAACCTATTAATTTCCCCTCGTCAAAAATAAGGTTATCAAGTGAGAAATCAC
CATGAGTGACGACTGAATCCGGTGAGAATGGCAAAAGTTTATGCATTTCTTTCCAGACTTG
TTCAACAGGCCAGCCATTACGCTCGTCATCAAAATCACTCGCATCAACCAAACCGTTATTCA
TTCGTGATTGCGCCTGAGCGAAACGAAATACGCGATCGCTGTTAAAAGGACAATTACAAAC
AGGAATCGAATGCAACCGGCGCAGGAACACTGCCAGCGCATCAACAATATTTTCACCTGAA
TCAGGATATTCTTCTAATACCTGGAATGCTGTTTTCCCAGGGATCGCAGTGGTGAGTAACC
ATGCATCATCAGGAGTACGGATAAAATGCTTGATGGTCGGAAGAGGCATAAATTCCGTCAG
CCAGTTTAGTCTGACCATCTCATCTGTAACATCATTGGCAACGCTACCTTTGCCATGTTTCA
GAAACAACTCTGGCGCATCGGGCTTCCCATACAATCGATAGATTGTCGCACCTGATTGCCC
GACATTATCGCGAGCCCATTTATACCCATATAAATCAGCATCCATGTTGGAATTTAATCGCG
GCCTAGAGCAAGACGTTTCCCGTTGAATATGGCTCATACTCTTCCTTTTTCAATATTATTGA
AGCATTTATCAGGGTTATTGTCTCATGAGCGGATACATATTTGAATGTATTTAGAAAAATAAA
CAAATAGGGGTTCCGCGCACATTTCCCCGAAAAGTGCCACCTGACGTCTAAGAAACCATTA
TTATCATGACATTAACCTATAAAAATAGGCGTATCACGAGGCCCTTTTGTC
